# “Plasma Membrane Calcium ATPase Downregulation in Dopaminergic Neurons Induces Presynaptic Dysfunction and Neuronal Vulnerability In Vivo and In Vitro”

**DOI:** 10.64898/2026.04.30.721667

**Authors:** Brenda Erhardt, Valentina Koltyk, María Isabel Farías, Mauricio Ricardo Bruno Dellepiane, Fernando Juan Pitossi, María Celeste Leal

## Abstract

Plasma Membrane Calcium ATPase (PMCA) is essential for maintaining intracellular calcium homeostasis. Previously, we used constitutive PMCA downregulation in *Drosophila melanogaster* dopaminergic neurons as a model to increase intracellular calcium and mimic early neuronal alterations associated with Parkinson’s disease. Here, we examined the mechanisms underlying the effects mediated by the conditional, adult-specific downregulation of PMCA in dopaminergic neurons in *Drosophila melanogaster*, both in vivo and in primary neuronal cultures. Adult-specific conditional silencing of PMCA in dopaminergic neurons reduced lifespan but to a lesser extent than the constitutive model and impaired locomotor performance. At the cellular level, PMCA-downregulated dopaminergic neurons exhibited elevated basal calcium, indicating disrupted calcium regulation. This was associated with a progressive increase in presynaptic vesicles and extracellular dopamine levels, suggesting enhanced neurotransmitter release. Notably, the synaptic active zone structure was preserved, indicating primarily functional rather than structural alterations. In primary neuronal cultures, PMCA downregulation reduced dopaminergic neuron survival and induced transient increases in neurite branching. Together, these findings show that PMCA downregulation leads to calcium dysregulation and presynaptic dysfunction without overt neurodegeneration in vivo, while promoting premature neuronal death in culture, indicating increased vulnerability and supporting a pre-degenerative state in which synaptic alterations precede neuronal loss.

## Introduction

To investigate the role of Plasma Membrane Calcium ATPase (PMCA) in dopaminergic neurons, we previously developed a model in *Drosophila melanogaster* in which PMCA was constitutively downregulated using RNA interference (PMCA^RNAi^) driven by the tyrosine hydroxylase (TH)-Gal4 driver (1, 2). This model exhibited reduced survival and elevated intracellular calcium (Ca^2+^) levels, leading to increased mitochondrial reactive oxygen species (ROS), synaptic vesicle accumulation, and dopamine levels, along with locomotor impairments in the absence of overt neurodegeneration. These findings suggest that PMCA downregulation disrupts Ca^2+^homeostasis and may contribute to neuronal dysfunction. Given the presence of dopaminergic deficits without detectable neurodegeneration, we hypothesized that this model reflects an early pre-degenerative stage and sought to further investigate synaptic function over time. However, its markedly reduced lifespan (∼13 days), likely due to *TH-Gal4* expression in the gut (2), limits the analysis of long-term progression.

To address this limitation, we developed complementary in vivo and in vitro approaches to study the progressive effects of PMCA downregulation and Ca^2+^dysregulation. Specifically, we generated an adult-specific PMCA knockdown model in dopaminergic neurons and used primary neuronal cultures with dopaminergic neurons expressing PMCA^RNAi^ to monitor progression at the single-cell level. Using these approaches, we established a pre-neurodegenerative model characterized by neuronal dysfunction in the absence of overt cell loss, likely underlying the observed motor deficits. In vitro, we developed a simplified system to monitor disease-relevant features over time, enabling the assessment of progressive changes associated with PMCA^RNAi^, including increased vulnerability and premature neuronal death.

## Materials and Methods

### *Drosophila* strains and maintenance

Fly stocks and crosses were raised on standard cornmeal-yeast-agar medium under a 12 h/12 h light/dark cycle in 4-inch plastic vials, and were transferred to fresh food every 2–4 days. For adult-specific expression, males carrying Tub-Gal80^ts^ were analyzed at 15, 20, and 25 days of age. To achieve adult-specific expression of PMCA^RNAi^, crosses were maintained in a room at 18 °C during development, and newly hatched flies were shifted to 28 °C in an incubator (I-30BLL, Percival) to induce PMCA^RNAi^ expression. All experiments were performed using male flies maintained at 28 °C (20 flies per vial). TH (pale)-Gal4 #8848, Tub-Gal80^ts^ #7017, UAS CaMPARI2 #78317, UAS mCD8:mCherry #27391, UAS Syt.eGFP #6925, UAS DA1m #80048 and UAS EGFP #6874 were obtained from Bloomington Stock Center; UAS PMCA^RNAi^ #101743 was obtained from Vienna *Drosophila* RNAi Center and UAS Brp.mRFP was kindly provided by Dr. Ceriani.

### Survival assay

Adults were collected within 24 h after hatching of the imago and a total of 84–268 males per experimental group were monitored for survival, evaluating the number of dead flies every 2 days. This data is presented in a curve showing the percentage of flies alive and the average lifespan of each genotype using the Kaplan-Meier approach, according to (3) with some modifications.

### Climbing assay (rapid iterative negative geotaxis, RING)

We conducted the experiment until approximately 200 flies per genotype had been quantified. For each assay, 20 flies were transferred into a 50 mL graduated cylinder in which a 9 cm height mark had been drawn. The cylinder was gently tapped against the bench in a controlled manner and the flies were allowed to rest for a few seconds. It was then tapped gently three additional times to bring all flies to the bottom. The flies’ behavior was recorded for 30 s. The percentage of flies that climbed above the 9 cm mark was quantified every 2 s over a total period of 20 s. This protocol was adapted from (4).

### Ca^2+^ imaging by CaMPARI2 reporter

To measure basal intracellular Ca^2+^ in the PPM3 dopaminergic neurons cluster, flies were anesthetized one by one on ice and their brains were dissected in adult hemolymph-like solution (AHL) (108 mM NaCl, 8.2 mM MgCl2, 4 mM NaHCO3, 1 mM NaH2PO4, 2 mM CaCl2, 5 mM KCl, 5 mM Hepes, 80 mM sucrose, pH 7.3). Brains were embedded in 1 μl of 1% low melt agarose (Biozym Scientific GmbH, Hessisch Oldendorf, Germany) with posterior side facing upwards and were then bathed in 10 μl of AHL. After localizing PPM3 neurons with a Zeiss LSM 710 NLO confocal microscope, they were illuminated with a 405 nm laser for 3 min, with full-field scanning at 1 s intervals, to induce CaMPARI2 photoconversion. CaMPARI signal was recorded using a 488 nm Argon laser and a 543 nm laser. Z-stacks images that included all PPM3 neurons were obtained with a 20x immersion objective (NA: 1; 15-20 Z-slices). Fluorescence intensity of 488 nm (CaMPARI2 in absence of Ca^2+^) and 543 nm (CaMPARI2 in presence of Ca^2+^) was measured from sum Z-projections in the PPM3 somas using the ImageJ program. The 543/488 nm fluorescence ratio was calculated to quantify basal Ca^2+^ levels.

### Dopaminergic vesicles measurement by Syt.eGFP reporter

Heads from 15 and 25-day-old flies were fixed for 25 min at RT in 4% paraformaldehyde in PB. Dissected brains were mounted in Mowiol. Z-stacks were taken using a Zeiss LSM 710 NLO confocal microscope. Syt.eGFP signal was recorded using a 488 nm Argon laser. Digital images including the protocerebral bridge (PB) and the fan shaped body (FSB) were obtained with a 20x immersion objective (NA: 1; 35-50 Z-slices). Fluorescence intensity was measured from sum Z-projections generated separately for the PB and FSB regions using the ImageJ program.

### Dopamine presence by DA1m reporter

To measure dopamine in the extracellular synaptic cleft in the PB, flies were anesthetized one by one and their brains were dissected in adult AHL. Brains were embedded in 1 μl of 1% low melt agarose with the posterior side facing upwards and were then bathed in 10 μl of AHL. Z-stacks were taken using a Zeiss LSM 710 NLO confocal microscope. DA1m signal was recorded using a 488 nm Argon laser. Digital images including the protocerebral bridge (PB) were obtained with a 20x immersion objective (NA: 1; 20-25 Z-slices). Fluorescence intensity was measured from sum Z-projections in the PB region using the ImageJ program.

### Synaptic active zones measurement by Brp.RFP reporter

Heads from 25-day-old flies were fixed for 25 min at RT in 4% paraformaldehyde in PB. Dissected brains were mounted in Mowiol. Z-stacks were taken using a Zeiss LSM 880 confocal microscope. Brp.RFP signal was recorded using a 543 nm laser. Digital images focused on the fan shaped body were obtained with a 40x water immersion objective (NA: 1.2; 22-37 Z-slices). Fluorescence intensity was measured from sum Z-projections in the FSB region using the ImageJ program.

### Primary neuronal culture

Primary neuronal cultures were obtained from third-instar wandering larvae of Drosophila melanogaster following an adapted protocol from (5). Larval brains were dissected in Leibovitz medium supplemented with 10% fetal bovine serum (FBS) and 1% antibiotics, then washed in fresh medium and transferred to Rinaldini buffer supplemented with 1% antibiotics, where they were kept on ice until processing was completed. Tissue dissociation was performed by incubation with Collagenase I (0.5 mg/ml) for 1 h at room temperature under agitation, followed by three washes and mechanical dissociation by ∼200 gentle pipetting steps. The resulting cell suspension was filtered through a 40 µm cell strainer, and cell viability was assessed by Trypan Blue exclusion using a Neubauer chamber. Cells were seeded onto Concanavalin A-coated glass-bottom dishes (15 μg/ml, Sigma-Aldrich, C5275) at a density of 400.000 cells per 35 mm plate, and maintained in Leibovitz medium supplemented with 10% FBS, 1% antibiotics, and 2.5 µg/ml Amphotericin B at 25 °C. The medium was half-replaced every three days, and cells were analyzed at 3, 6, and 9 days post-seeding. Cells were seeded on Concanavalin A-coated (Sigma-Aldrich, C5275) dishes at a density of 400,000 cells per 35 mm glass-bottom plate, and maintained in Leibovitz medium supplemented with 10% FBS, 1% antibiotic and 2.5 µg/ml Amphotericin B at 25°C. Medium was half-changed every three days, and cells were analyzed at days 3, 6 and 9 post-seeding.

For coating, dishes were prepared the day prior to culture by incubation with 15 μg/ml Concanavalin A at 37 °C for 2 h, followed by a wash with apyrogenic water. Dishes were then allowed to dry overnight at 37 °C before use.

### Survival analyses in primary neuronal culture

To assess the survival of dopaminergic neurons expressing PMCA^RNAi^, primary cultures were generated from *TH-Gal4*>EGFP/PMCA^RNAi^ and *TH-Gal4*>EGFP (control) larvae. At 3 days in vitro (DIV), Z-stacks were taken using a Zeiss LSM 880 Airyscan inverted confocal microscope. EGFP+ neurons were recorded using a 488 nm Argon laser. Digital images including neuron’s somas were obtained with a 40x water immersion objective (NA: 1.2; 15-25 Z-slices). The same neurons were re-imaged at 6 and 9 DIV. To enable longitudinal tracking, dishes were temporarily sealed with parafilm during imaging to minimize contamination and returned to the incubator afterward. A reference mark was made on the bottom of each dish, defined as (0,0) coordinates, allowing relocation of the same fields at later time points using image metadata.

Neuronal survival was quantified by measuring EGFP fluorescence intensity from sum Z-projections in somas over time, and neurons were classified as dead when their fluorescence dropped below one-third of their initial intensity at 3 DIV. This approach accounts for variability in baseline fluorescence between neurons by normalizing each cell to its own initial signal (also seen in dopaminergic neurons in ex vivo or fixed brains). The number of surviving neurons was quantified at 6 and 9 DIV relative to 3 DIV.

### Neurite outgrowth by Sholl analysis in primary neuronal culture

Sum Z-projections of EGFP+ neurons were generated for Sholl analysis. An intensity threshold was applied using the Otsu method, and neuronal arborization was analyzed using the Sholl Analysis plugin, with concentric circles at 5 μm intervals centered on the soma and extending beyond the longest neurite. Neurite intersections at each radius were manually quantified using the Cell Counter plugin in ImageJ.

### Statistical analysis

Analyses were performed with GraphPad Prism 8.0. Outliers were identified and excluded using the ROUT method with Q = 1. Normality distribution was assessed using Shapiro–Wilk test and Kolmogorov–Smirnov test. When data were normally distributed, it was analysed using unpaired t test, one sample t test or one-way ANOVA with Tukey’s multiple comparisons test and results are represented as interleaved scatter with bars plot (mean SEM). When data were not normal, it was analysed with Mann–Whitney test or Kruskal–Wallis test with Dunn’s multiple comparisons test and results are represented with box and whiskers plot (median, quartile 1 and quartile 2). Two-way ANOVA was performed with Tukey’s multiple comparisons test or Sidak’s multiple comparisons test and results were represented as mean SEM. The tests used in each figure are detailed in figure legends. Statistical significance level was α = 0.05.

## Results

### The adult-specific expression of PMCA^RNAi^ under the *TH-Gal4* driver decreased fly lifespan

To understand the effect of increased Ca^2+^ in dopaminergic neurons in adult individuals and avoiding possible developmental effects, we expressed the PMCA^RNAi^ in dopaminergic neurons (*TH-Gal4* driver) only in adulthood using the system Tubulin-Gal80^ts^ (Tub-Gal80 thermosensitive). Once flies hatch, they are transferred to an incubator at 28°C to allow PMCA^RNAi^ expression. We previously reported that flies in the constitutive model *TH-Gal4*>PMCA^RNAi^ died earlier compared to control (13 days vs. 44 days) (2). Here we investigate if adult-specific PMCA^RNAi^ expression in dopaminergic neurons had a longer lifespan than the constitutive model. We found that flies *TH-Gal4*,Tub-Gal80^ts^>PMCA^RNAi^ displayed a shorter lifespan than control genotypes *TH-Gal4*,Tub-Gal80^ts^/+ and UAS PMCA^RNAi^/+ (30 days vs. 40-44 days, **Figure S1**), possibly due to *TH-Gal4* expression in the gut (2). In summary, we established a model in which PMCA downregulation in dopaminergic neurons reduces mean lifespan, while extending the temporal window for assessing dopaminergic phenotypes, as flies survive longer than in the constitutive model.

**Figure S1.**
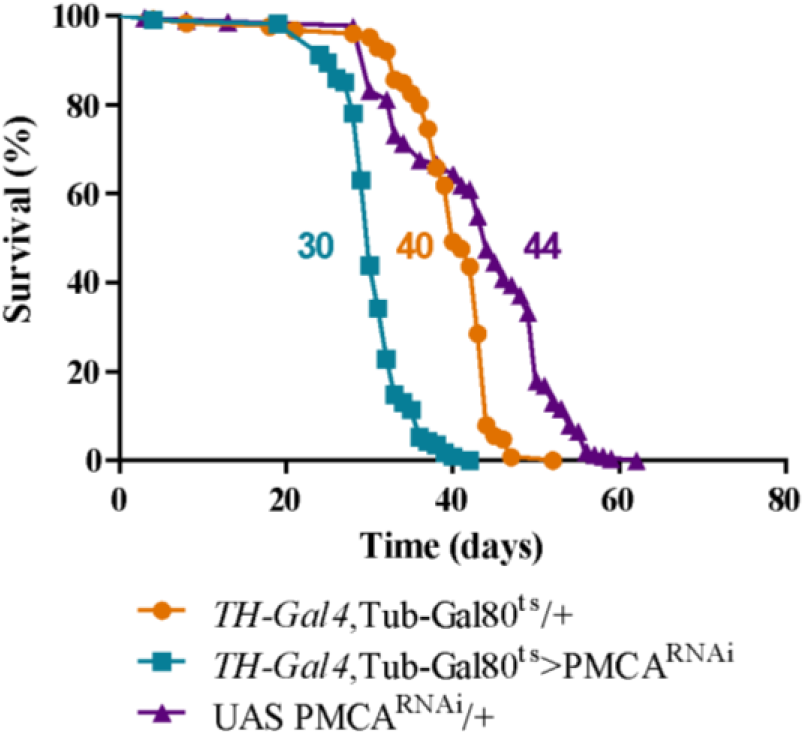
Adult-specific expression of PMCA^RNAi^ under *TH-Gal4* driver reduced flies’ lifespan. Survival curves show the percentage of flies alive per day. PMCA^RNAi^-expressing flies exhibit a shorter lifespan compared to controls at 28°C. The number next to each curve indicates the mean lifespan of each genotype. Curves were analyzed using the log-rank (Mantel–Cox) test. Each condition was replicated three times (n = 114–213 flies).

### Adult-specific PMCA^RNAi^ expression in TH neurons impaired locomotion

We next assessed whether adult-specific PMCA^RNAi^ expression in dopaminergic neurons induces motor alterations over time. We measured climbing performance at 15, 20 and 25 days post-hatch. We analyzed the percentage of flies that climbed above 9 cm every 2s over a 20s period. *TH-Gal4*,Tub-Gal80^ts^>PMCA^RNAi^ flies climbed less than controls (*TH-Gal4*,Tub-Gal80^ts^ and UAS-PMCA^RNAi^/+) at 15 and 25 days. At 20 days, *TH-Gal4*,Tub-Gal80^ts^>PMCA^RNAi^ flies climbed less than *TH-Gal4*,Tub-Gal80^ts^ but did not differ significantly from UAS-PMCA^RNAi^/+, which showed a reduced locomotor activity compared to their performance at 15 days (**Figure 1**). In *Drosophila*, age-associated locomotor decline is well documented (6). As presented in Figure 1, there is a general decline when comparing 15 to 25 days. The decreased climbing phenotype in the control genotype UAS-PMCA^RNAi^/+ prompts us to speculate decreased fitness in some situations, as we previously reported in (1). Overall, these results indicate that adult-specific PMCA downregulation in dopaminergic neurons leads to a progressive impairment in locomotor activity.

**Figure 1.**
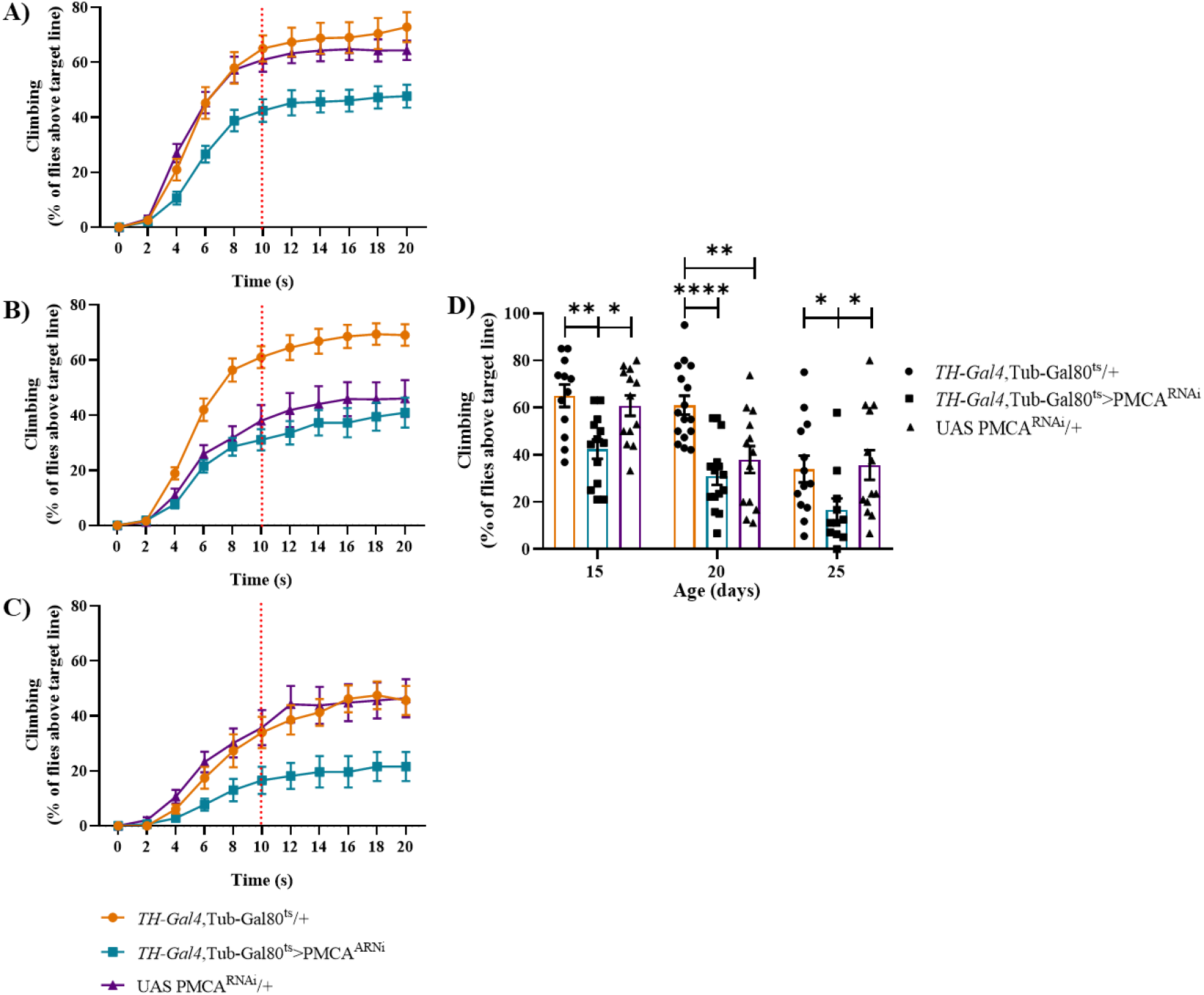
Adult-specific expression of PMCA^RNAi^ in dopaminergic neurons impaired climbing ability. (A–C) Percentage of flies that climbed above 9 cm every 2 s over a 20 s period at 15 (A), 20 (B), and 25 (C) days of age. The red dotted line indicates the time point used for the data shown in (D). (D) Percentage of flies that exceeded 9 cm after 10 s at 15, 20, and 25 days of age. (D) Two-way ANOVA with Tukey’s multiple comparisons test (n = 11–16): at 15 days, *TH-Gal4*,Tub-Gal80^ts^/+ vs. *TH-Gal4*,Tub-Gal80^ts^>PMCA^RNAi^, P = 0.0052; *TH-Gal4*,Tub-Gal80^ts^>PMCA^RNAi^ vs. UAS-PMCA^RNAi^/+, P = 0.0242; at 20 days, *TH-Gal4*,Tub-Gal80^ts^/+ vs. *TH-Gal4*,Tub-Gal80^ts^>PMCA^RNAi^, P < 0.0001; *TH-Gal4*, Tub-Gal80^ts^/+ vs. UAS-PMCA^RNAi^/+, P = 0.002; at 25 days, *TH-Gal4*,Tub-Gal80^ts^/+ vs. *TH-Gal4*,Tub-Gal80^ts^>PMCA^RNAi^, P = 0.0468; *TH-Gal4*, Tub-Gal80^ts^>PMCA^RNAi^ vs. UAS-PMCA^RNAi^/+, P = 0.0252.

### Adult-specific expression of PMCA^RNAi^ under *TH-Gal4* increased intracellular Ca^2+^levels in PPM3 somas

We next investigated whether PMCA downregulation increases basal intracellular Ca^2+^. To address this, we used the CaMPARI2 sensor to measure basal Ca^2+^ in PPM3 neurons. In our previous work, the constitutive model showed elevated Ca^2+^ in the same neuronal cluster where the TH driver was expressed, which corresponds to the dopaminergic population analogous to the human *substantia nigra*, a region implicated in PD (7). Here the PPM3 neurons of *TH-Gal4*,Tub-Gal80^ts^>CAMPARI2/PMCA^RNAi^ flies exhibited higher Ca^2+^ levels compared to *TH-Gal4*,Tub-Gal80^ts^>CAMPARI2 control, with 174% vs. 93% of the 543/488 nm fluorescence ratio, respectively (**Figure 2**). This ratio serves as a proxy for intracellular Ca^2+^ levels. Measurements were restricted to 15-day-old flies, as the combination of CaMPARI2 expression and PMCA^RNAi^ appeared to reduce viability, suggesting potential toxicity of the reporter gene under these conditions.

**Figure 2.**
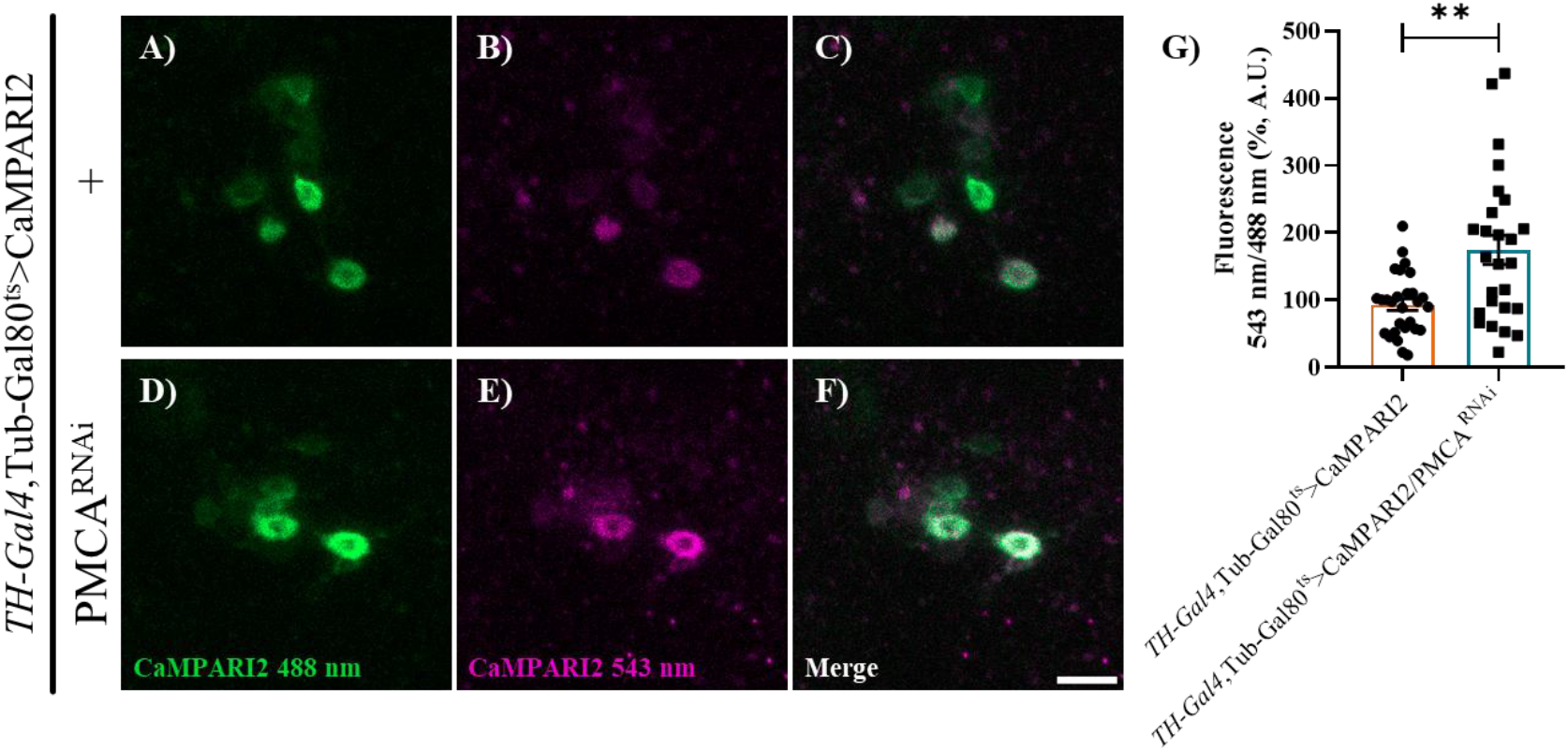
Adult-specific PMCA^RNAi^ expression in TH neurons increased basal Ca^2+^ levels at 15 days old flies. Basal Ca^2+^ was assessed by CaMPARI2 fluorescence in PPM3 somas. (A–F) Representative images of PPM3 somas after photoconversion with 405 nm light in flies expressing CaMPARI2 under the *TH-Gal4* driver (A–C) or co-expressing CaMPARI2 and PMCA^RNAi^ (D–F). Following photoconversion, CaMPARI2 fluoresces at 488 nm in the absence of Ca^2+^ (A, D) and at 543 nm (shown in magenta) in the presence of Ca^2+^ (B, E). (C, F) Merged images showing the relative contribution of green and red signals; colocalization appears white. (G) Quantification of the 543 nm/488 nm CaMPARI2 fluorescence ratio reveals increased cytosolic Ca^2+^ in adult-specific PMCA^RNAi^-expressing neurons compared to control. Data were normalized to the median of the control group within each experiment (A.U.: arbitrary units). Scale bar: 10 μm. Statistical analysis was performed using Mann Whitney test (n =26–28); P = 0.0033.

### Adult-specific expression of PMCA^RNAi^ under *TH-Gal4* led to a progressive increase in presynaptic vesicles

Even though flies expressing PMCA^RNAi^ in dopaminergic neurons under the TH driver exhibited reduced survival, impaired climbing ability, and elevated cytosolic Ca^2+^ in dopaminergic soma, we observed no evidence of neurodegeneration (data not shown). Therefore, we ask if these anomalies have a correlation with problems in dopaminergic synapsis that could explain the motor impairment observed. Our question is also supported by the fact that in our previous model of constitutive expression of PMCA^RNAi^ in dopaminergic neurons, we found increased levels of synaptic vesicles and dopamine in the presynapsis (1 and data not shown, respectively). To find out if the dopaminergic synapses were altered over time, we analyzed the presynaptic terminals in the Protocerebral Bridge (PB) and Fan Shape Body (FSB) (regions that have been proposed to be analogous to the human striatum, the projection target of *substantia nigra* dopaminergic neurons, which is affected in Parkinson’s disease (7)) in fixed-brains of 15 and 25 days-old. We used the Syt.eGFP reporter to estimate synaptic vesicle abundance, with eGFP fluorescence serving as a proxy for vesicle presence. We found that synaptic vesicle levels were increased in dopaminergic presynaptic terminals of the PB and FSB in flies expressing PMCA^RNAi^, compared to mCherry-expressing controls (**Figure 3**). Moreover, in PMCA^RNAi^-expressing flies, vesicle accumulation increased over time, as indicated by higher eGFP fluorescence intensity at 25 days compared to 15 days. Together, these results suggest that synaptic function in the central complex is disrupted upon PMCA downregulation.

**Figure 3.**
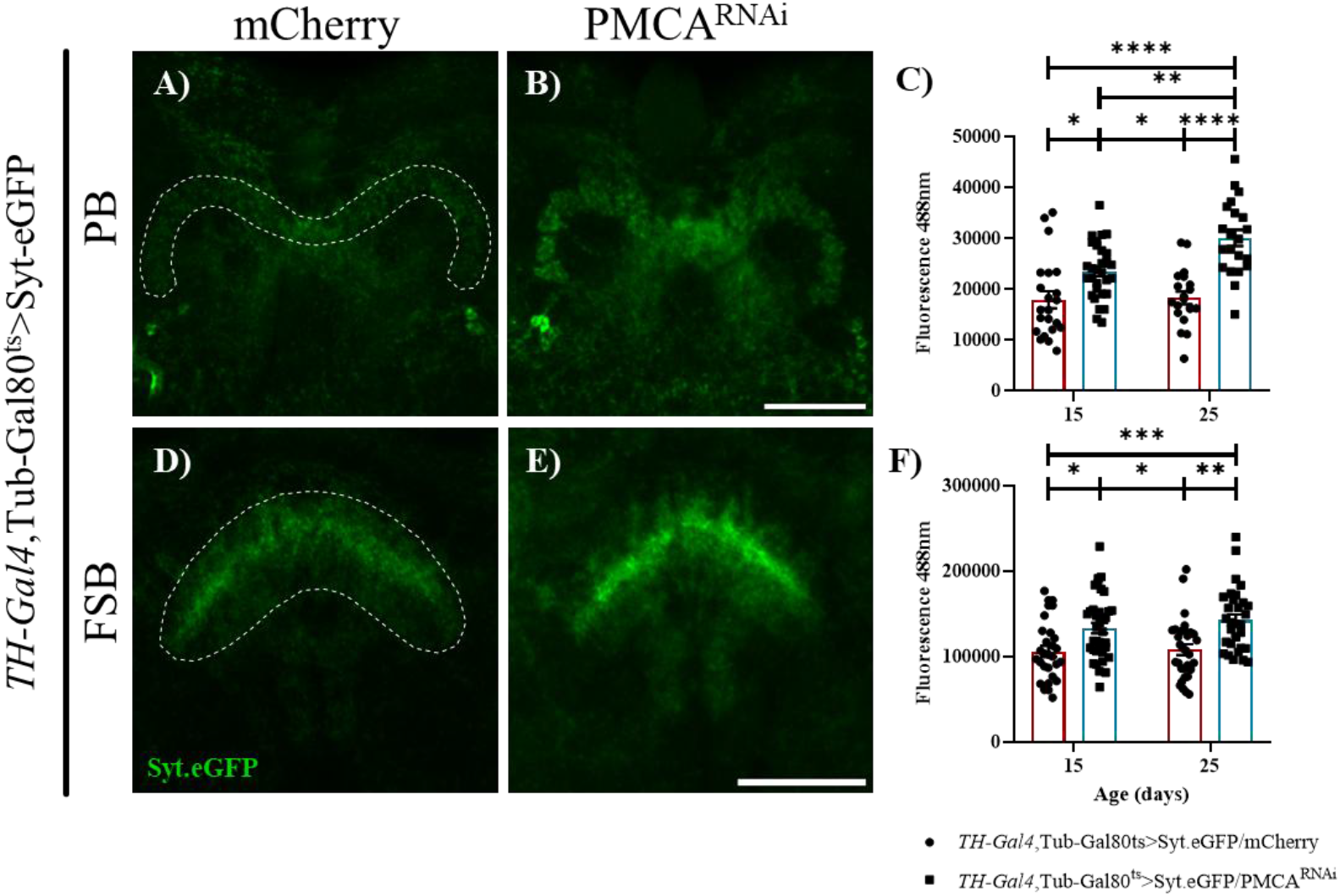
Adult-specific expression of PMCA^RNAi^ under *TH-Gal4* generated a progressive increase in synaptic vesicles in the PB and FSB. Synaptic vesicle levels were assessed by Syt.eGFP fluorescence in dopaminergic projections. (A-B, D-E) Representative images of the PB and FSB, respectively, in flies with adult-specific expression of Syt.eGFP under the *TH-Gal4* (A, D) or co-expressing Syt.eGFP and PMCA^RNAi^ (B, E) at 25 days. (C, F) Quantification of Syt.eGFP fluorescence intensity in the PB (C) and FSB (F) shows a progressive increase in synaptic vesicle levels in PMCA^RNAi^-expressing flies compared to controls. Scale bar: 50 μm. Statistical analysis was performed using two-way ANOVA followed by Tukey’s multiple comparisons test (C, n = 20-30 and F, n = 31-37); *p < 0.05, **p < 0.01, ***p < 0.001, and ****p < 0.0001.

### Adult-specific expression of PMCA^RNAi^ under *TH-Gal4* produced a progressive increase in extracellular dopamine level in the presynapsis

We next asked whether the increase in synaptic vesicles also augmented neurotransmitter levels. To determine this, we measured dopamine at 15 and 25 days using the extracellular dopamine reporter DA1m in the PB. Due to *ex vivo* imaging conditions, increased tissue thickness limited optical access to deeper brain regions; therefore, the FSB could not be reliably imaged, and analysis was restricted to the PB. We found that extracellular dopamine was incremented compared to control and showed a progressive rise over time (**Figure 4**).

**Figure 4.**
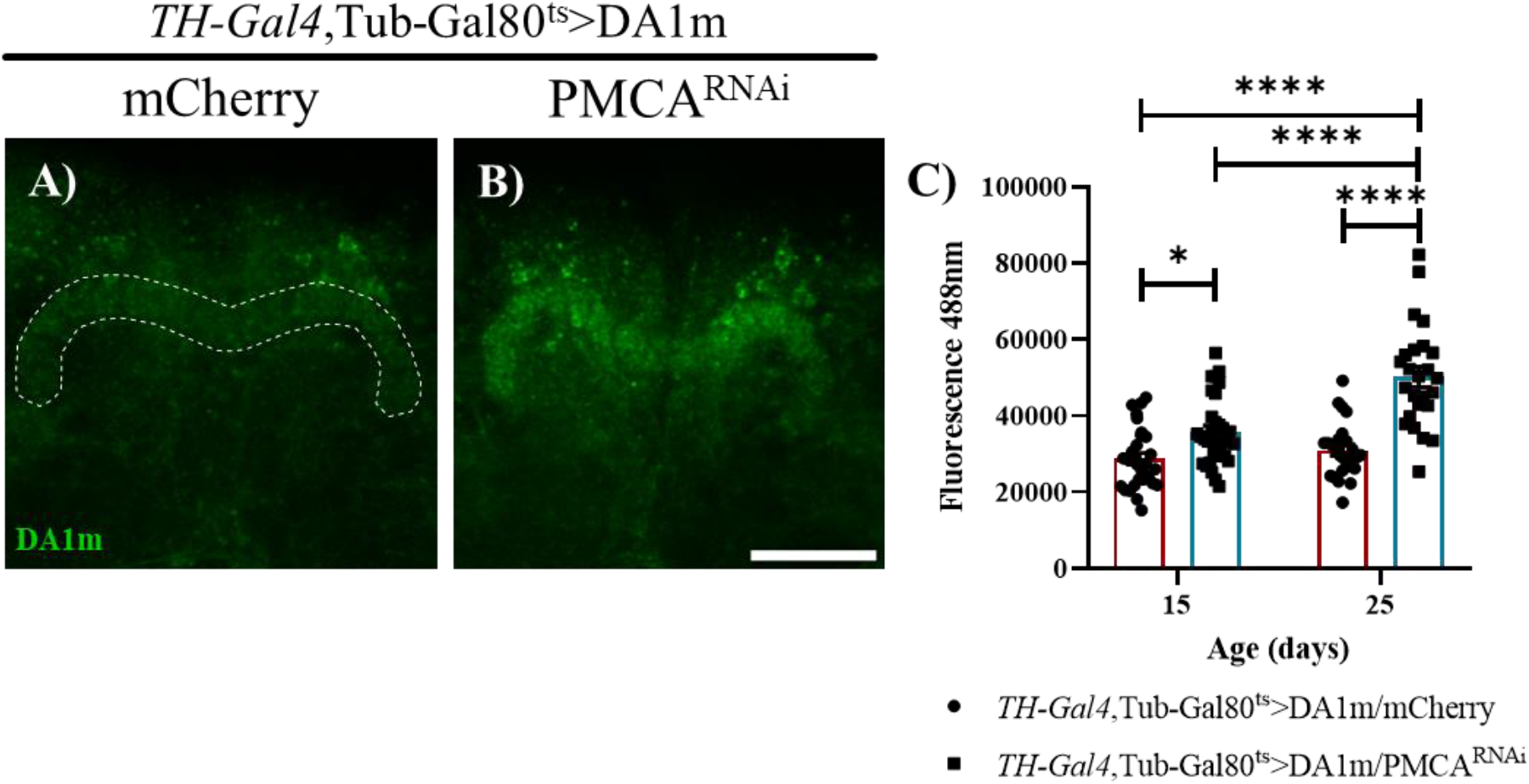
Adult-specific expression of PMCA^RNAi^ under *TH-Gal4* led to a progressive increase in extracellular dopamine level in the PB. Extracellular dopamine was assessed using the DA1m sensor in dopaminergic projections. (A– B) Representative images of the protocerebral bridge (PB) in flies with adult-specific co-expression of DA1m and mCherry under *TH-Gal4* (A) or co-expression of DA1m and PMCA^RNAi^ (B) at 25 days. (C) Quantification of DA1m fluorescence intensity in the PB shows increased extracellular dopamine levels in PMCA^RNAi^-expressing flies compared to controls. Scale bar: 50 μm. Statistical analysis was performed using two-way ANOVA followed by Tukey’s multiple comparisons test (n = 25-32); *p < 0.05, and ****p < 0.0001.

### Specific expression of *TH-Gal4*,Tub-Gal80^ts^>PMCA^RNAi^ did not produce an increase in presynaptic BRP levels

Given the observed increase in presynaptic vesicles and dopamine levels, we next assessed whether synaptic structural organization was altered. To this end, we measured Bruchpilot (BRP) levels at 25 days, using the Brp.mRFP reporter, which labels active presynaptic zones, in the FSB. We compared brains from *TH-Gal4*, Tub-Gal80^ts^>EGFP/PMCA^RNAi^;Brp.mRFP flies with control flies (*TH-Gal4*, Tub-Gal80^ts^>EGFP;Brp.mRFP) (**Figure 5**). We found no significant differences in Brp.mRFP fluorescence intensity between genotypes, indicating that the structural organization of presynaptic active zones was not affected by PMCA^RNAi^.

**Figure 5.**
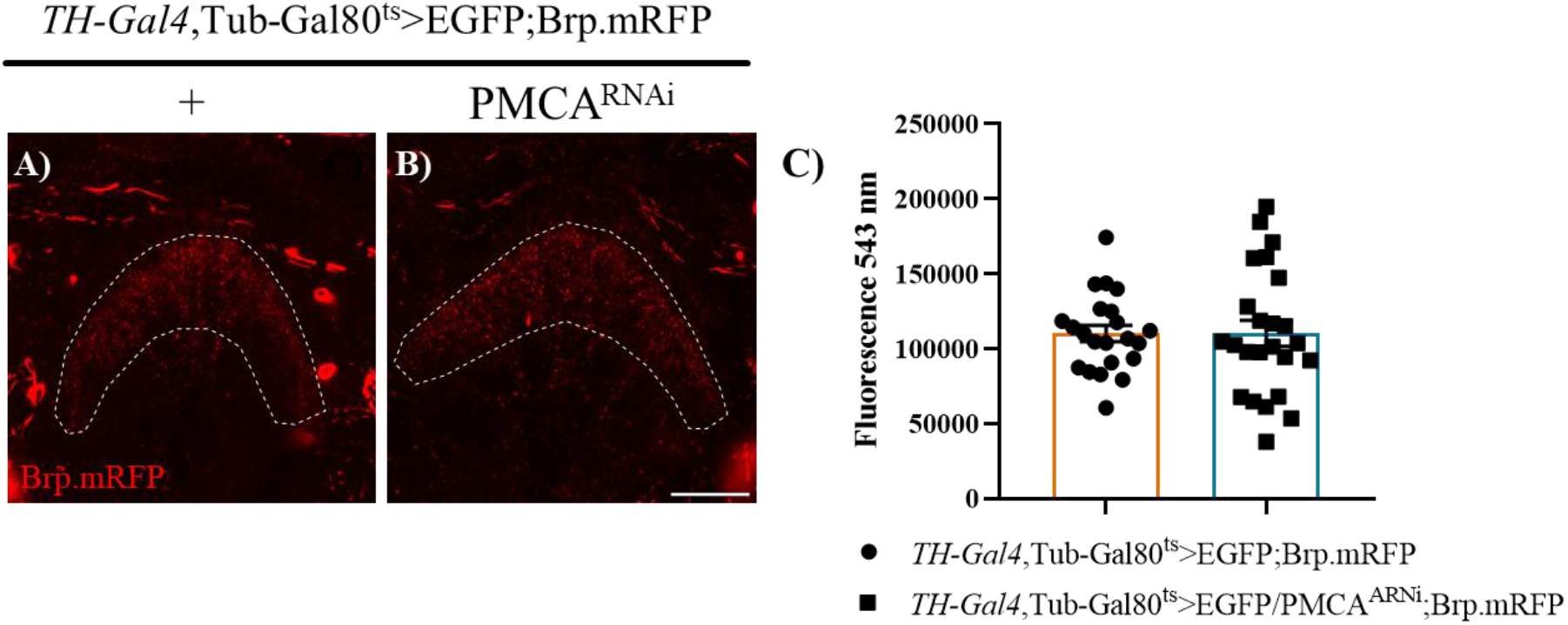
Adult-specific expression of PMCA^RNAi^ under *TH-Gal4* did not produce alteration in presynapsis Brp.RFP+ in the FSB. Measurement of BRP was assessed using Brp.mRFP sensor in dopaminergic terminals. (A–B) Representative images of the FSB in flies with adult-specific co-expression of EGFP and Brp.mRFP under *TH-Gal4* (A) or co-expression of EGFP, PMCA^RNAi^ and Brp.mRFP (B) at 25 days. (C) Quantification of Brp.mRFP fluorescence intensity in the FSB did not show differences in BRP expression between PMCA^RNAi^-expressing flies compared to controls. Scale bar: 25 μm. Statistical analysis was performed using unpaired t-test (n = 21–24); P=0,9922.

### Dopaminergic neurons expressing PMCA^RNAi^ exhibited reduced survival at 9 days *in vitro*

To further investigate the progression of these phenotypes at the single-cell level, we turned to primary neuronal cultures expressing PMCA^RNAi^ in dopaminergic neurons (under *TH-Gal4* driver), allowing longitudinal tracking of individual cells. To this end, we established cultures from larval brains in which dopaminergic neurons co-expressed PMCA^RNAi^ and EGFP for tracking, or EGFP alone in controls, and assessed neuronal survival at 3, 6, and 9 days in vitro (DIV), as well as neuronal morphology. As shown in **Figure 6** dopaminergic neurons co-expressing PMCA^RNAi^ and EGFP exhibited reduced survival compared to control neurons expressing EGFP alone, indicating that downregulation of PMCA in dopaminergic neurons in culture are more vulnerable than controls.

**Figure 6.**
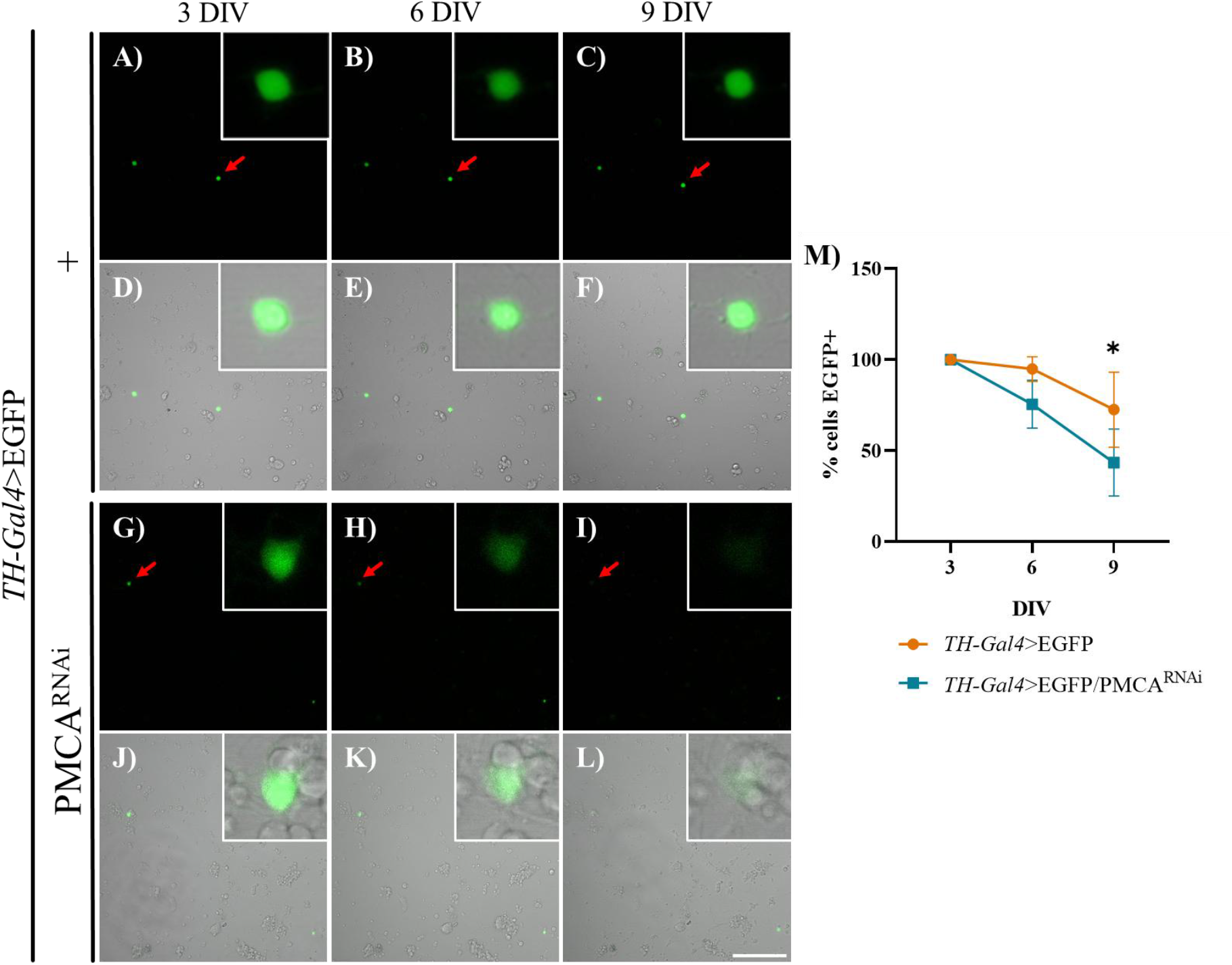
Dopaminergic neurons expressing PMCA^RNAi^ exhibited reduced survival at 9 days *in vitro*. (A–L) Representative images of *TH-Gal4*>EGFP neurons (A–F) and *TH-Gal4*>EGFP/PMCA^RNAi^ neurons (G–L). EGFP+ neurons at 3, 6, and 9 DIV (A–C, G–I) and merged images with the corresponding brightfield channels (D–F, J–L). Red arrows indicate the neurons shown in the magnified images. (M) Percentage of EGFP+ neurons in control and PMCA^RNAi^ cultures, expressed relative to the number of cells at 3 days in vitro (DIV), with longitudinal tracking of the same cells at 6 and 9 DIV shows that PMCA^RNAi^-expressing dopaminergic neurons had lower survival than control cells at 9 DIV. Scale bar: 100 μm. Statistical analysis was performed using two-way ANOVA followed by Sidak’s multiple comparisons test (n = 4 independent cultures); *p < 0.05.

### Neurons expressing PMCA^RNAi^ under *TH-Gal4* showed increased neurite branching near the soma at 3 and 6 days *in vitro*

To assess whether PMCA^RNAi^-expressing dopaminergic neurons display morphological alterations, we performed Sholl analysis of neuronal structure at 3, 6, and 9 DIV. At 3 DIV, dopaminergic neurons with PMCA downregulation (TH-Gal4>EGFP/PMCA^RNAi^) showed a significantly higher number of intersections from neurites into Sholl circles compared to control neurons (TH-Gal4>EGFP), indicating a broader, more elaborate branching pattern concentrated near the soma (∼10–70 μm) (**Figure 7**C). This increase in neurite complexity was maintained, although attenuated, at 6 DIV, where significant differences between genotypes were restricted to intersections closer to the soma (∼10–15 μm and ∼50 μm) (**Figure 7**D). By 9 DIV, no significant differences in the number of intersections were detected between PMCA^RNAi^ and control neurons at any distance from the soma (**Figure 7**E). Together, these results indicate that PMCA downregulation induces an early, transient increase in dendritic branching complexity near the soma, an effect that progressively diminishes and is no longer detectable at later stages of neuronal maturation in culture.

**Figure 7.**
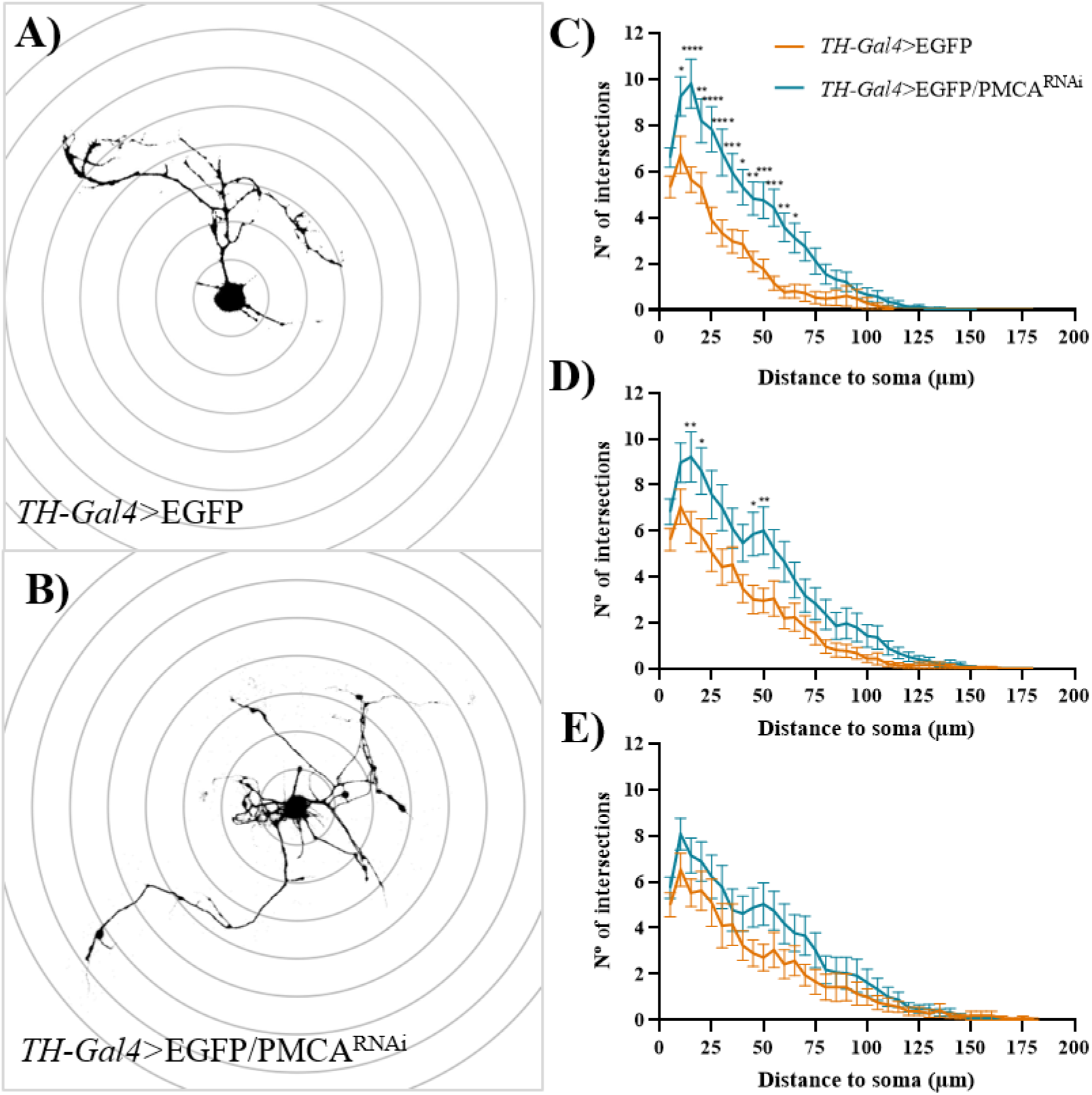
PMCA^RNAi^ expression in dopaminergic neurons altered neuronal morphology. Sholl analysis was performed on EGFP+ neurons. (A–B) Representative images of control neurons at 3 DIV (A) and neurons expressing PMCA^RNAi^ (B). (C–E) Number of neurite intersections at increasing distances from the soma (every 5 μm) at 3 (C), 6 (D), and 9 DIV (E). PMCA^RNAi^-expressing neurons show altered neurite complexity compared to controls. Statistical analysis was performed using two-way ANOVA followed by Sidak’s multiple comparisons test. 3 DIV, n= 60 neurons; 6 DIV, n= 49 neurons; 9 DIV, n= 48 neurons. *p< 0.05, **p< 0.01, ***p< 0.001, ****p< 0.0001; *TH-Gal4*>EGFP/PMCA^RNAi^ vs. *TH-Gal4*>EGFP.

## Discussion

Here, we developed an adult-specific model of PMCA^RNAi^ in dopaminergic neurons in Drosophila. This model leads to increased basal intracellular Ca^2+^levels, altered presynaptic function, and locomotor deficits in the absence of dopaminergic neuron loss. Compared to the constitutive model (1), the phenotype is milder; however, the extended lifespan allows for a more detailed analysis of disease progression.

The adult-specific PMCA^RNAi^ model exhibited impaired climbing performance at all ages analyzed. This is partially consistent with our previous findings in the constitutive PMCA^RNAi^ model (1), although the time points examined differ between studies. As expected, an age-dependent decline in climbing ability was also observed, consistent with previous reports (8).

Importantly, although no neurodegeneration was observed in vivo, dopaminergic neurons exhibited reduced survival in vitro, suggesting that PMCA^RNAi^ induces a state of cellular vulnerability that may be context-dependent. *In vitro*, PMCA^RNAi^-expressing dopaminergic neurons displayed increased neurite arborization near the soma. These changes in neurite complexity, potentially associated with alterations in cytoskeletal organization, resemble those reported in amyloid *in vitro* models (9). PMCA^RNAi^ may promote alterations in the filamentous actin structure, potentially contributing to the branching phenotype observed (10). Future experiments will further address this possibility. Whether this structural remodeling contributes to the reduced survival observed in these neurons remains to be determined.

Taken together, our results suggest that PMCA downregulation in TH-positive neurons increases dopamine release and disrupts synaptic function while increasing neuronal vulnerability, without causing overt neuronal loss. These findings indicate that dopaminergic neuron loss is not required to elicit the locomotor deficits and altered dopamine signaling observed in *Drosophila*. Consistent with this, a parkin knockdown model also displays alterations in synaptic activity in the absence of clear neurodegeneration (11).

More broadly, our data support a model in which early neuronal dysfunction precedes and potentially predisposes to degeneration, rather than being a mere consequence of cell loss. In this context, PMCA downregulation–induced Ca^2+^dysregulation may act as a vulnerability factor that sensitizes neurons to additional stressors, aligning with the concept of Parkinson’s disease as a “multiple-hit” process, where neuronal impairment and degeneration emerge from the interaction between intrinsic vulnerability and subsequent insults (12).

## Acknowledgments

We thank Dr. Eduardo Miguel Castaño for lending fly materials, Dr. María Fernanda Ceriani and her group as well as Dr. Pablo Wappner and his group from IIBBA— Leloir Institute for providing some fly lines and technical assistance. We thank Andrés Liceri for the fly food, Dr. Andrés H. Rossi and Dr. Alejandra Ross from the Microscopy and Imaging Facility at the Leloir Institute Foundation for his support and assistance. This work was supported by grants of René Baron Foundation to Fernando J. Pitossi and Brenda Erhardt, PICT2020-56-ANPCyT and Young IBRO Maternity to M. Celeste Leal.

## Bibliography

1- Erhardt B, Marcora MS, Frenkel L, Bochicchio PA, Bodin DH, Silva BA, Farías MI, Allo MÁ, Höcht C, Ferrari CC, Pitossi FJ, Leal MC. Plasma membrane calcium ATPase downregulation in dopaminergic neurons alters cellular physiology and motor behaviour in Drosophila melanogaster. Eur J Neurosci. 2021 Sep;54(6):5915–5931. doi: 10.1111/ejn.15401. Epub 2021 Aug 16. PMID: 34312939.

2- Erhardt B, Frenkel L, Marcora MS, Farías MI, Ferrari CC, Pitossi FJ, Leal MC. Downregulation of Plasma Membrane Ca2+ ATPase driven by tyrosine hydroxylase-Gal4 reduces Drosophila lifespan independently of effects in neurons. Fly (Austin). 2023 Dec;17(1):2192457. doi: 10.1080/19336934.2023.2192457. PMID: 36949021; PMCID: PMC10038040.

3- Linford NJ, Bilgir C, Jennifer R, et al. Measurement of lifespan in drosophila melanogaster. J Visualized Exp. 2013;JoVE(71). DOI: 10.3791/50068

4- Madabattula ST, Strautman JC, Bysice AM, O’Sullivan JA, Androschuk A, Rosenfelt C, Doucet K, Rouleau G, Bolduc F. Quantitative Analysis of Climbing Defects in a Drosophila Model of Neurodegenerative Disorders. J Vis Exp. 2015 Jun 13;(100):e52741. doi: 10.3791/52741. PMID: 26132637; PMCID: PMC4544889.

5- Egger B, van Giesen L, Moraru M, Sprecher SG. In vitro imaging of primary neural cell culture from Drosophila. Nat Protoc. 2013 May;8(5):958–65. doi: 10.1038/nprot.2013.052. Epub 2013 Apr 18. PMID: 23598446.

6- Jones MA, Grotewiel M. Drosophila as a model for age-related impairment in locomotor and other behaviors. Exp Gerontol. 2011 May;46(5):320–5. doi: 10.1016/j.exger.2010.08.012. Epub 2010 Aug 26. PMID: 20800672; PMCID: PMC3021004.

7- Nicholas J. Strausfeld,Frank Hirth. Deep Homology of Arthropod Central Complex and Vertebrate Basal Ganglia.Science340,157-161(2013). DOI:10.1126/science.1231828

8- Privalova V, Szlachcic E, Sobczyk Ł, Szabla N, Czarnoleski M. Oxygen Dependence of Flight Performance in Ageing Drosophila melanogaster. Biology (Basel). 2021 Apr 14;10(4):327. doi: 10.3390/biology10040327. PMID: 33919761; PMCID: PMC8070683.

9- Saad Y, Segal D, Ayali A. Enhanced neurite outgrowth and branching precede increased amyloid-β-induced neuronal apoptosis in a novel Alzheimer’s disease model. J Alzheimers Dis. 2015;43(3):993–1006. doi: 10.3233/JAD-140009. PMID: 25125474.

10- Dalghi MG, Ferreira-Gomes M, Rossi JP. Regulation of the Plasma Membrane Calcium ATPases by the actin cytoskeleton. Biochem Biophys Res Commun. 2018 Nov 25;506(2):347–354. doi: 10.1016/j.bbrc.2017.11.151. Epub 2017 Nov 24. PMID: 29180009.

11- Eduard Dumitrescu, Jeffrey M Copeland, B Jill Venton “Parkin Knockdown Modulates Dopamine Release in the Central Complex, but Not the Mushroom Body Heel, of Aging Drosophila”. ACS Chem Neurosci 2023 Jan 18;14(2):198–208. doi: 10.1021/acschemneuro.2c00277. Epub 2022 Dec 28., PMID: 36576890, PMCID: PMC9897283. DOI: 10.1021/acschemneuro.2c00277

12- Leal MC, Casabona JC, Puntel M, Pitossi FJ. Interleukin-1β and tumor necrosis factor-α: reliable targets for protective therapies in Parkinson’s Disease? Front Cell Neurosci. 2013 Apr 29;7:53. doi: 10.3389/fncel.2013.00053. PMID: 23641196; PMCID: PMC3638129.

